# Functional principal component based time-series genome-wide association in sorghum

**DOI:** 10.1101/2020.02.16.951467

**Authors:** Chenyong Miao, Yuhang Xu, Sanzhen Liu, Patrick S. Schnable, James C. Schnable

## Abstract

The phenotypes of plants develop over time and change in response to the environment. New engineering and computer vision technologies track phenotypic change over time. Identifying genetic loci regulating differences in the pattern of phenotypic change remains challenging. In this study we used functional principal component analysis (FPCA) to achieve this aim. Time-series phenotype data was collected from a sorghum diversity panel using a number of technologies including RGB and hyperspectral imaging. Imaging lasted for thirty-seven days centered on reproductive transition. A new higher density SNP set was generated for the same population. Several genes known to controlling trait variation in sorghum have been cloned and characterized. These genes were not confidently identified in genome-wide association analyses at single time points. However, FPCA successfully identified the same known and characterized genes. FPCA analyses partitioned the role these genes play in controlling phenotype. Partitioning was consistent with the known molecular function of the individual cloned genes. FPCA-based genome-wide association studies can enable robust time-series mapping analyses in a wide range of contexts. Time-series analysis can increase the accuracy and power of quantitative genetic analyses.

## Introduction

Quantitative genetic approaches are widely used across all domains of biology to identify genetic loci controlling variation in target traits. Many genes where naturally occurring functionally variable alleles control variation in agronomically relevant traits have been identified in crops using both QTL mapping (structured populations) or genome-wide association studies (association panels)^1–3^. Collected phenotypic data from the hundreds to thousands of accessions required for QTL mapping or genome-wide association study (GWAS) has historically been a time and resource intensive undertaking. Hence most attempts to identify genes controlling variation in a target trait employ data from a single time point, usually either at maturity or a fixed number of days after planting. However recent engineering – wearable devices, automated phenotyping greenhouses, field phenotyping robots, and unmanned aerial vehicles (UAVs) – and computer vision advances are lowering the barriers and activation energy requires to score traits from multiple points throughout development^4–11^. Plant growth and development is a dynamic process, responding to environmental perturbations and regulated by different suites of genes at different times and in different environments^4, 10–15^. The availability and use of time-series trait data from mapping and association populations has the potential to increase both the accuracy and power of gene mapping studies^4, 16^. It may also provide greater insight into the biologically distinct roles different loci pay in determining final phenotypes. However, integrating time-series data into statistical frameworks originally envisioned for single trait measurements across large populations is not straightforward. A range of differing approaches are currently being explored by the community.

There are several approaches to the use of time-series phenotypic data in mapping studies. The most straightforward is to conduct QTL mapping or GWAS separately at each time point and then summarize the mapping results^12, 17, 18^. However, this approach is not robust to missing data or subsets of the population being scored on alternating time points and requires complex approaches to multiple testing correction given the partially correlated nature of both linked genetic markers and measurements of the same trait in the same individuals at multiple time points. A second widely employed method is to summarize patterns of change over time using pre-defined functions with discrete numbers of variables^19–21^. QTL mapping or GWAS can then be conducted for the values of the different variables within the equation as that function is fit to observed data from different individuals. This approach can be powerful and is able to impute missing data points in individuals. However, this approach can fail when the pattern of phenotypic change over time follows an unknown function, too complex to fit, or does not conform to the expected function fit the observations^11^. Functional principal component analysis (FPCA) as a more general method provides some of the strengths of fitting parametric functions – sharing data across time points, ability to impute missing values – without requiring patterns of phenotypic change over time fit any particular function^22–24^. A variation of nonparametric functional principal component based mapping has been successfully employed to identify loci controlling the gravitropism response in *A. thaliana* seedlings^4, 22^. Kwak and coworkers concluded that dimensional reduction via FPCA may increase power to detect QTL in recombinant inbred populations relative to prior approaches that include trait data from each time point^22^. Muraya included an FPCA method adapted from Kwak^22^ to identify several trait associated SNPs for maize biomass accumulation in vegetative development. This represented an advance over parametric regression, which had not been able to identify any trait associated SNPs using the same trait and marker dataset. However, none of the identified loci coincided with either markers identified in single time point analyses or known loci controlling the trait of interest.^15^.

Here we employ a new approach to functional principal component analysis of non-parametric regression data^23^ and evaluate its effectiveness at both controlling false positives and identifying known true positives in sorghum. We employ a phenotypically constrained subset of the sorghum association panel (SAP)^25^. This population was grown and imaged through vegetative and reproductive development in a high throughput phenotyping facility. Organ level semantic segmentation from hyperspectral images was employed to extract phenotypic values^26^. Three cloned sorghum genes controlling height – *dwarf1, dwarf2*, and *dwarf3* – are segregating in the Sorghum Association Panel. *Dwarf1* (*dw1*, Sobic.009g229800) belongs to a previously uncharacterized protein family in plants has been shown to influences plant height in sorghum, rice, arabidopsis by reducing cell proliferation activity in the internodes and appears to act in the brassinosteroid signaling pathway^27–29^. *Dwarf2* (*dw2*, Sobic.006G067700) encodes a protein kinase^30^. *Dwarf3* (*dw3*, Sobic.007G163800) encodes a MDR transporter orthologous to *brachytic2* in maize and appears to influence cell elongation through a role in polar auxin transport^31^. We demonstrate that genome-wide association using the functional principal component scores derived from the approach described in by Xu and co-workers^23^ can be used to successfully identify all three known true positive genes. Three novel signals were also detected, one of which can be independently validated in data from study of a different sorghum population. These results contrast favorably terminal phenotyping or time point by time point analyses in the same population, and to provide additional insight into the distinct biological roles each of the three known genes play in determining sorghum height.

## Results

### Loss of association power in phenotypically constrained populations

The classical sorghum dwarfing genes have large effects on plant height and are segregating within the SAP^32, 33^. Field collected plant height data for 357 lines from the SAP correctly identified both *dw1* and *dw2* [Figure 1A] as well as a third previously reported locus known to influence variation in height in this population^32, 33^. Thirty-eight lines from the field study exceeded the physical limit on maximum height for the imaging facility employed in this study [Figure 1B]. After the exclusion of these 38 lines as well as an additional 27 lines which failed to germinate or failed to thrive in the greenhouse, field measured height data for the remaining 292 lines was still sufficient to identify *dw1* – though with substantially reduced statistical significance – however, after the removal of these phenotypically extreme lines, signals from *dw2* and the other previously reported height controlling loci were no longer statistically significant [Figure 1C]. *Dwarf3*, located on the long arm of chromosome 7, was not identified in association analyses using either field data on terminal plant heights from complete set of 357 SAP lines, or in analyses of data only from the phenotypically constrained subset of SAP lines employed for greenhouse experiments.

**Figure 1.**
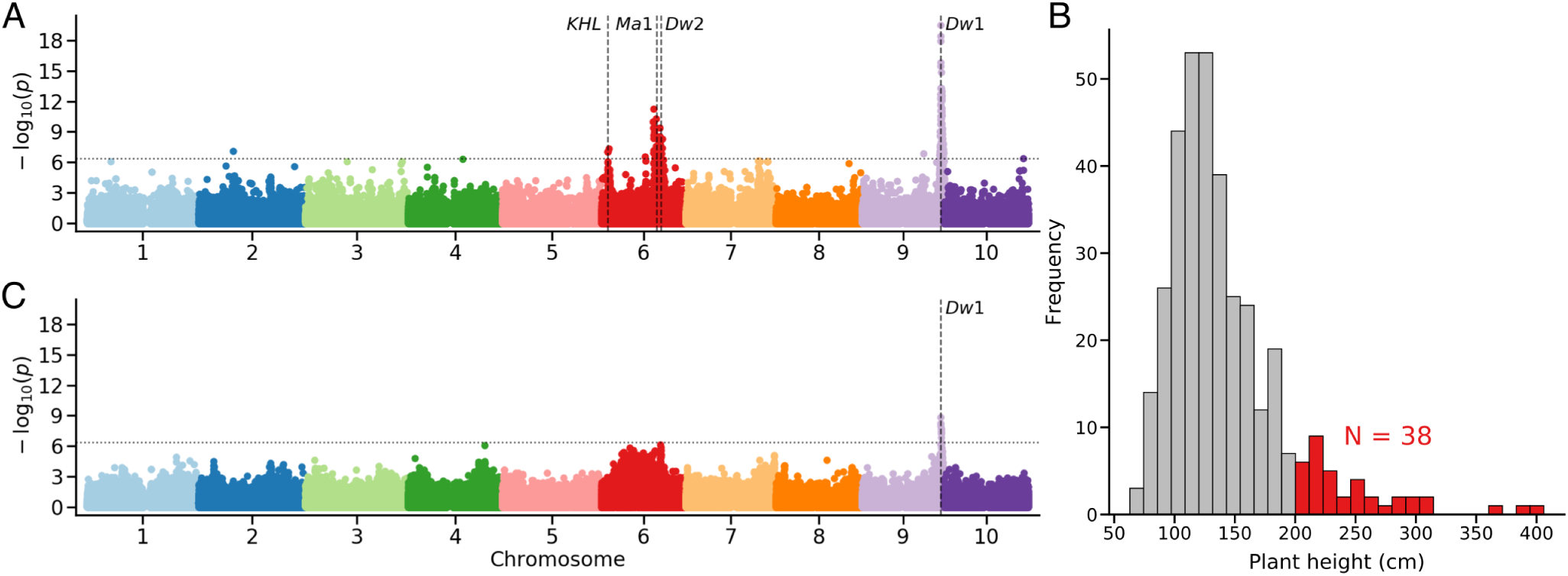
Reduced power to identify causal loci in phenotypically constrained populations. (A) A genome wide association analyses for plant height, defined as the distance between the soil surface and the top of the panicle at maturity, using field collected data for 357 lines from sorghum association panel and the set of genotype call data used in this study. The location of *dw1, dw2* and *ma1* are indicated with dashed lines, as is an additional known height locus (KHL) identified in multiple prior GWAS conducted on height in this population using different genetic marker data. (B) Distribution of observed heights for the 357 lines employed for association analysis in panel A. The set of thirty eight lines above two meters in height are marked in red. (C) An genome wide association analysis identical to that shown in panel A but with the exclusion of lines with heights >2 meters (38 lines) and those which we were not able to successfully germinate and phenotype in this study (27 lines).

### Genetic associations with sorghum height at different time points

The phenotypically constrained subset of 292 SAP lines were imaged at the University of Nebraska’s Greenhouse Innovation Center (UNGIC). Imaging lasted approximately one month centered on reproductive development and each plant was imaged using a set of five cameras including top views and side views from multiple angles and hyperspectral imaging from a single side view perspective^26, 34, 35^. Two methods were employed to extract plant height at individual time points. The first was whole plant segmentation, an approach widely used in plant image analysis^36^: segmentation of an image into plant pixels and not plant pixels, and measuring height by the difference between the minimum and maximum y-axis values for plant pixels [Figure 2A]. The second approach was a recently described method^26^ to semantically segment “plant pixels” of sorghum plants into separate stalk, leaf, and panicle classes [Figure S1, 2B]. There are many different ways to measure the plant height^26, 37^. Generally, researchers in the field have preferred to benchmark on the height of stalk, height to the uppermost leaf collar, or the height to the tip of the inflorescence rather than the height to the highest leaf tip. An advantage of the latter method – semantic segmentation – is that it can be used to measure a version of plant height which more closely approximates how height is measured in the field^26^. Sorghum plant height measured by whole plant segmentation tended to oscillate over time as new leaves emerged, while sorghum plant height measured via the semantic segmentation method tended to be monotonically increasing [Figure 2C]. Generally, researchers in the field have preferred to measure the height of stalk or the height of inflorescence rather than the height to the highest point on the plant, which is often a leaf tip. The semantic segmentation approach makes it possible measure height using a definition for plant height the corresponds to the definition of plant height used by sorghum geneticists in the field^26^.

**Figure 2.**
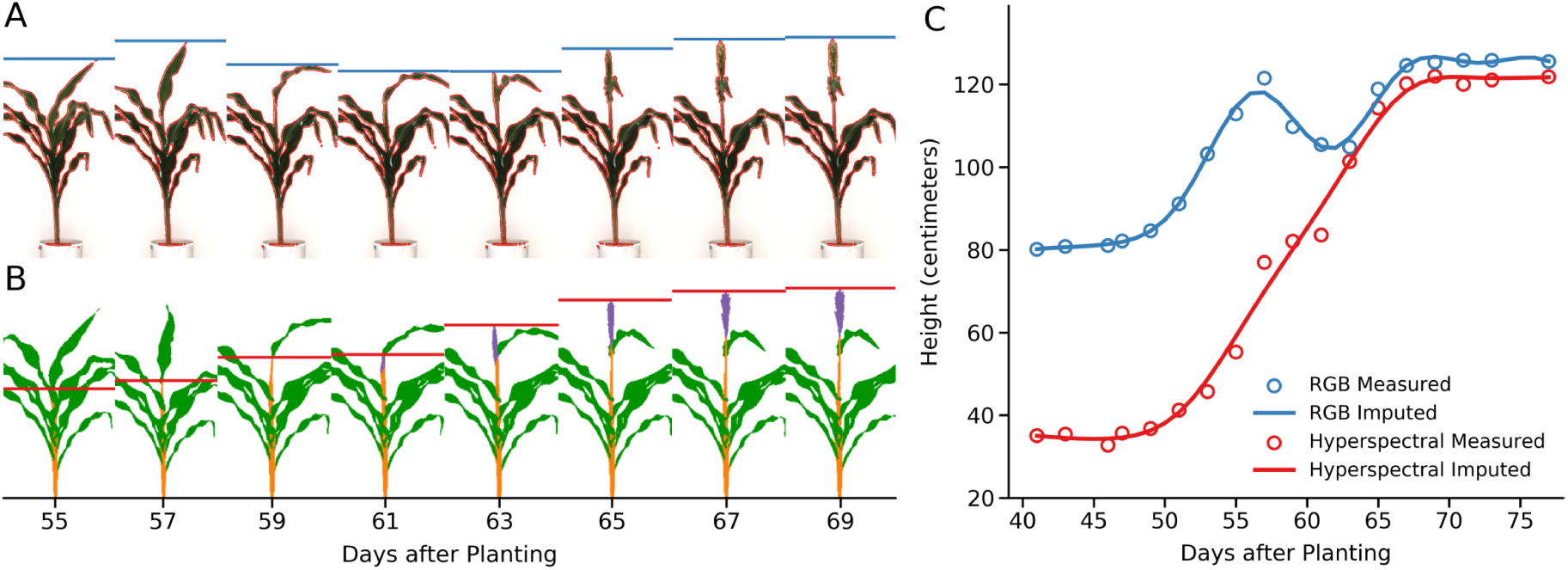
Different methods to define and measure plant height produce different outcomes. (A) Conventional RGB images of a single sorghum plant (PI 576401) taken on eight different days spanning the transition from vegetative to reproductive development. Pixels identified as “plant” through whole plant segmentation are outlined in red. Measured plant height, defined as the distance between the plant pixels with the smallest and greatest y-axis value is indicated by the horizontal blue bar in each image. (B) Semantically segmented images of same plant taken on the same day from a moderately different viewing angle using a hyperspectral camera. Pixels classified as “leaf” are indicated in green, pixels classified as “stem” are indicated in orange, and pixels classified as “panicle” are indicated in purple. Measured plant height, defined as the distance between stem or panicle pixels with the smallest and greatest y-axis value is indicated by the horizontal red bar in each image. (C) Observed and imputed plant heights for the same sorghum plant on each day within the range of phenotypic data collection. Blue and red circles indicate measured height values from whole plant segmentation of RGB images and semantic segmentation of hyperspectral images, respectively. Solid blue and solid red lines indicate height values imputed for unobserved time points using nonparametric regression for whole plant and semantic height datasets, respectively.

A second challenge faced by many time-series imaging studies is the maximum daily throughput provided by fixed imaging infrastructure. In this particular study, individual plants were divided into two groups imaged on alternating days. Therefore there was no single time point at which image or height data was available for all individual plants [Figure S2A]. In order to estimate the height of each individual plant at each individual time point, nonparametric regression was employed to impute all missing data points [Figure S2B]. To assess the accuracy of this imputation, the percent of variance in height explained by genetic factors was assessed at individual time points for subsets of the population using either height measured directly from images or height imputed from nonparametric curves. In most cases proportion of variance in imputed height which could be explained by genetic factors matched or exceeded the equivalent value for directly measured height [Figure S3]. This is consistent with previous work which found missing values in time-series plant phenomics data can be imputed accurately from nonparametric curves^24^.

Independent genome-wide association analyses were conducted for imputed sorghum height – as measured via semantic segmentation – on each day from 41 to 76 days after planting (DAP). Consistent with our observations from field collected plant height data, almost no significant trait/marker associations were observed in the analyses for DAP 65-76 [Figure S5]. On early days in the experiment statistically significant signals were identified all over the genome likely as a result of extreme height values for a small number of lines which had experienced reproductive transition and panicle emergence prior to the start of imaging [Figure S4]. Excluding extreme lines led to the identification of signals near *dw1, dw3*, and *ma3* in data from some early time points mixed in among approximately a dozen other repeatedly identified loci across the genome. A loss of detectable genetic associations between days 54 and 66 after planting corresponded roughly to booting and panicle exertion in many sorghum accessions. The peduncle length is under the control of a distinct set of genetic factors from plant height below the flag leaf^33^, and the timing of panicle exertion is under the control of a third distinct set of factors, maturity genes^38, 39^. We speculate that, once a large proportion of lines advanced to the booting stage or beyond, the role these three sets of genetic factors played in determining plant height diluted the total variance attributable to any one single genetic locus, reducing power to identify statistically significant associations.

A second set of analyses were conducted where nonparametric curves calculated for individual plants were aligned based on time relative to panicle emergence (e.g Days After Panicle Emergence or DAPE) rather than time relative to planting (Days After Planting or DAP) (Figure 3). Given variation in the date of panicle emergence and the dangers of extrapolating nonparametric curves beyond the range of observed data points it must be noted that this approach meant that data were available only for distinct subsets of the original 292 phenotyped sorghum lines at any given time point. Day by day genome-wide association studies were conducted from 14 days prior to panicle emergence to 10 days after panicle emergence, as described above. No known sorghum flowering time loci were identified using this approach, with the exception of *ma1*. However due to the strong linkage between *ma1* and *dw2* genes based on previous studies we cannot exclude the possibility that the significantly associated markers near *ma1* reflect the effect of *dw2* on plant height^40, 41^. The signal corresponding to *dw2* was identified in the DAPE GWAS analysis for six days between -14 to -5 DAPE with the signal 323 kilobases from the known locus. The signal corresponding to *dw1* was identified from -14 to +1 DAPE and was located only 15 kilobases away from the known location of *dw1* on sorghum chromosome 9. This was closer than the 114 kilobase distance between the gene and the closest significant SNP identified in DAP GWAS results [Figure S5]. *Dwarf3* did not show any statistically significant signal in the DAPE based analysis but was identified in some time points when plant data was compared using DAP. Hence, while it was possible to identify all three known true-positive genes controlling sorghum plant height through genome-wide association analyses at individual time points, in no case were all three identified in a single analyses, across the approximately sixty total genome-wide association studies conducted at different time points or using different developmental landmarks. Many other confounding associations were also identified. In many cases both the single most significantly associated SNP and “hot zone” of SNPs all showing strong significant association with trait variation was quite distant from the known causal locus although still within the range expected given observed LD decay rates in sorghum^32, 42^

**Figure 3.**
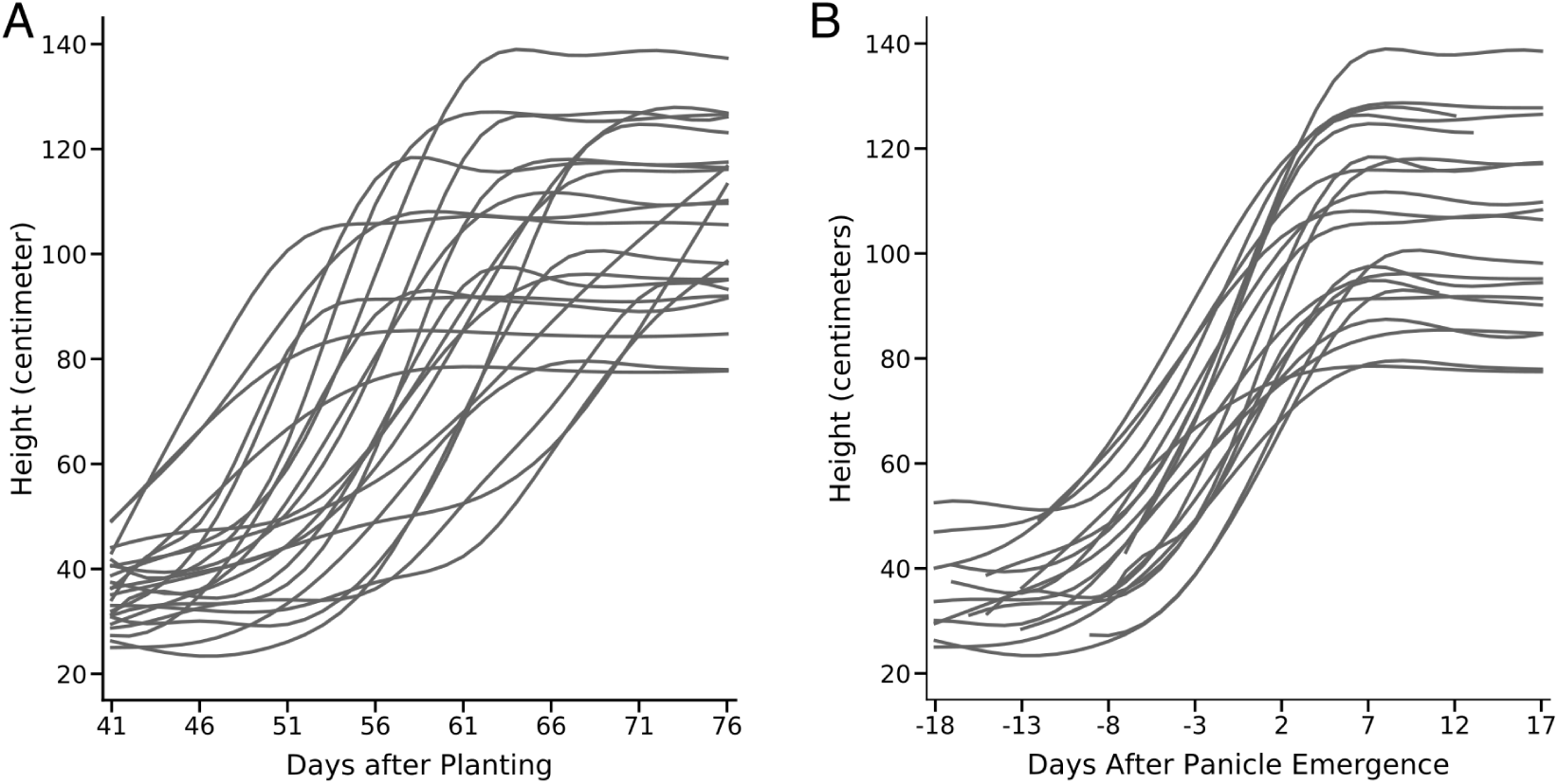
Comparison of change in plant height over time for members of the SAP population when anchoring either on planting date (DAP) or panicle emergence date (DAPE). (A) Growth curves imputed using nonparametric regression for 23 sorghum genotypes, anchored for comparison based on sharing the same date of planting. (B) Growth curves imputed using nonparametric regression for same 23 sorghum genotypes shown in panel A, anchored for comparison based on sharing the same date of panicle emergence.

**Figure 4.**
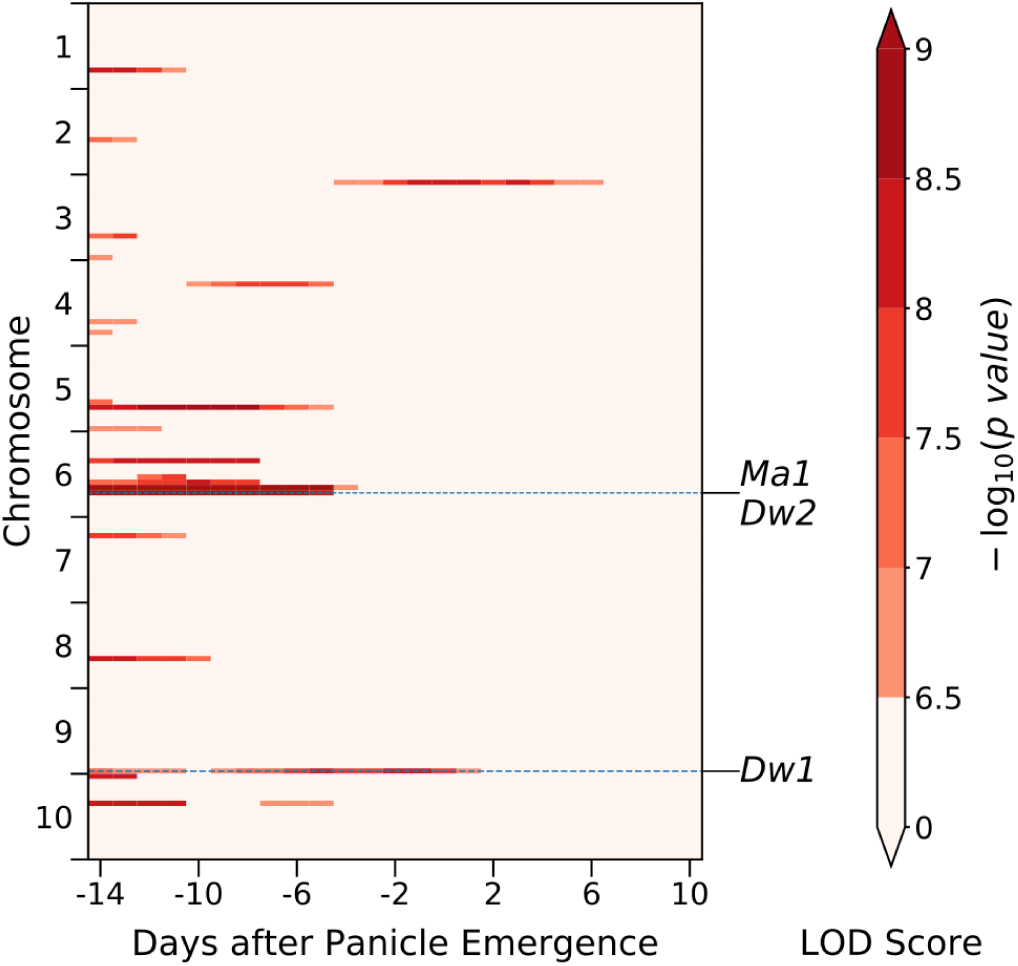
Several known causal loci show statistically significant associations with plant height when sequential genome wide association studies are conducted using data anchored to the date of panicle emergence. A summary of where statistically significant trait associated SNPs were identified in separate genome wide association studies conducted using height data for each day between -14 to +10 days after panicle emergence (DAPE). Each vertical column summarizes the results from one of the twenty five independently conducted genome wide association studies. Each sorghum chromosome is divided into sixteen bins containing equal numbers of SNP markers. Each cell in each vertical column is color coded based on the single most significant p-value observed for any marker within that bin on that day. Light pink cells indicate bins which contain no markers which exceed the multiple testing corrected threshold for statistical significance. The locations of the two cloned dwarf genes and the one cloned maturity gene which were successfully identified in analysis of data from at least one time point are indicated with horizontal dashed lines.

### Mapping genes controlling variation in growth curves

All three known true positive height genes were identified by sequential time point based GWAS. However, there was no single time point, nor in any single treatment of the data (DAP or DAPE) were all three ground truth genes were identified. Many other loci not previously reported to be linked to height were also identified with equal or greater statistical support to known true positive genes across many time points. In order to conduct the individual time point analyses, it was necessary to fit nonparametric curves to each individual plant to impute unobserved values. Functional principal component analyses [Figure 5A] was employed to decompose variation among the curves into four functional principal components that combined to explain >99% of the total variance in curve shape among the plants in the population. The first two functional principal components were able to explain >97% of total variation [Figure 5B & C].

**Figure 5.**
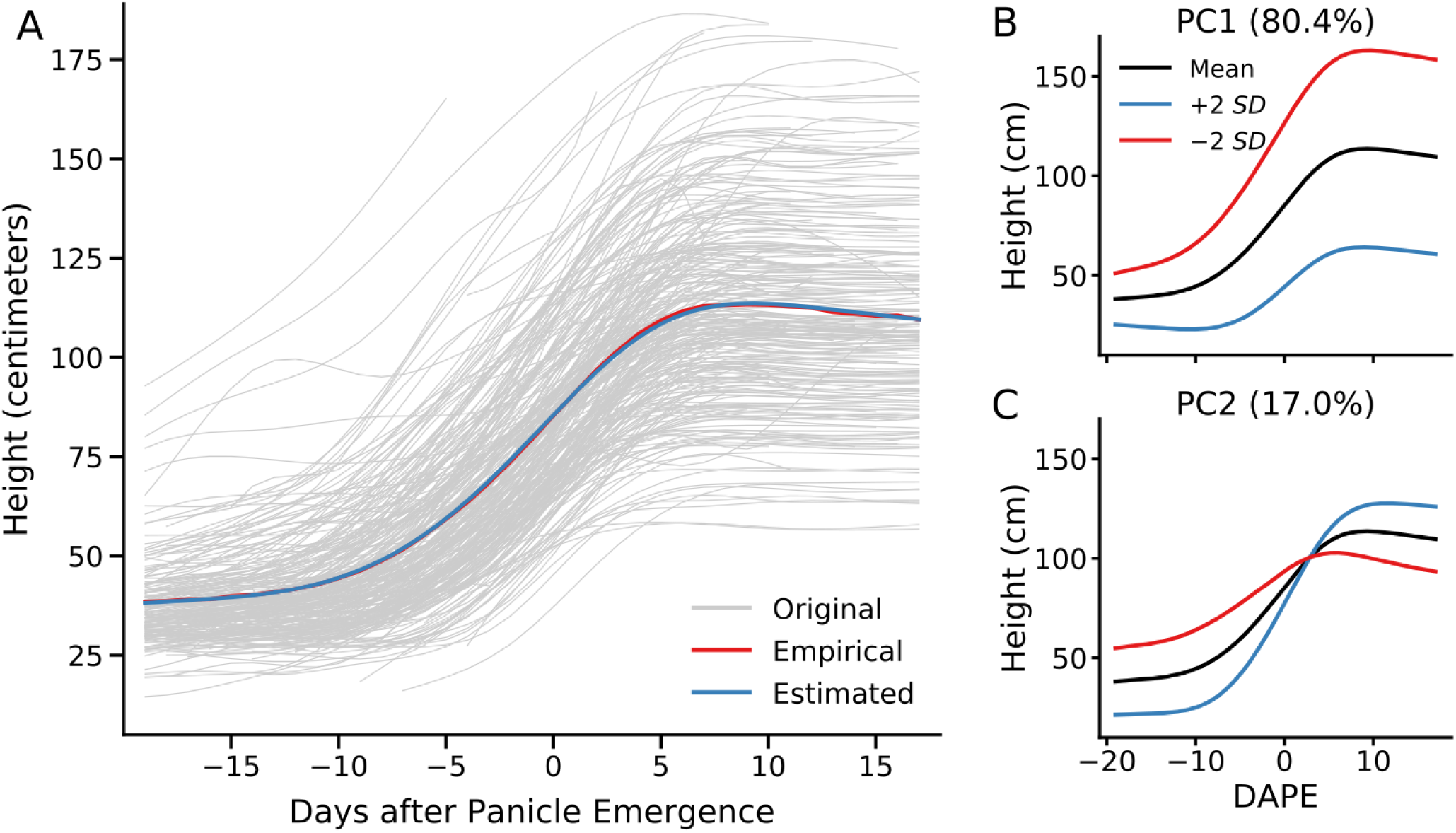
Two functional principal components explain >97% of variance in the sorghum growth curves observed in this study. Functional principal component analyses seeks to describe the pattern of change in height over time of each observed plant using a mean function combined with variable weightings of a set of eigenfunctions. (A) Comparison of the empirical mean function (red) for all growth curves observed in this study (gray) and the mean function estimated using functional principal component analysis (blue). (B) Illustration of how changing the score for functional principal component one alters the resulting growth curve. (SD = Standard Deviation) (C) Illustration of how changing the score for functional principal component two alters the resulting growth curve.

As the first two functional principal components described the vast majority of the variation in patterns of change in plant growth over time, genome-wide association analyses were conducted to identify genes controlling variation in these two phenotypic descriptors [Figure 6]. Sorghum lines with negative scores of the first functional principal component tended to be taller overall and exhibited greater increases in height during panicle emergence than lines with positive scores of the second functional principal component [Figure 6A & B]. Both *dw1* and *dw2* showed statistically significant associations with variation in the first functional principal component, as did a single locus on the short arm of chromosome three [Figure 6E]. Sorghum lines with negative functional principal component two scores tended to be taller prior to panicle emergence but exhibit limited additional increases in height during panicle emergence, while lines with positive functional principal component two scores started out shorter, but exhibited big increases in height during panicle emergence so that this second set of lines was taller at the end of the experiment [Figure 6C & D]. *Dwarf3* showed a statistically significant association with variation in the second functional principal component, as did two other regions of the genome close to the centromeres of sorghum chromosomes 5 and 9 [Figure 6F].

**Figure 6.**
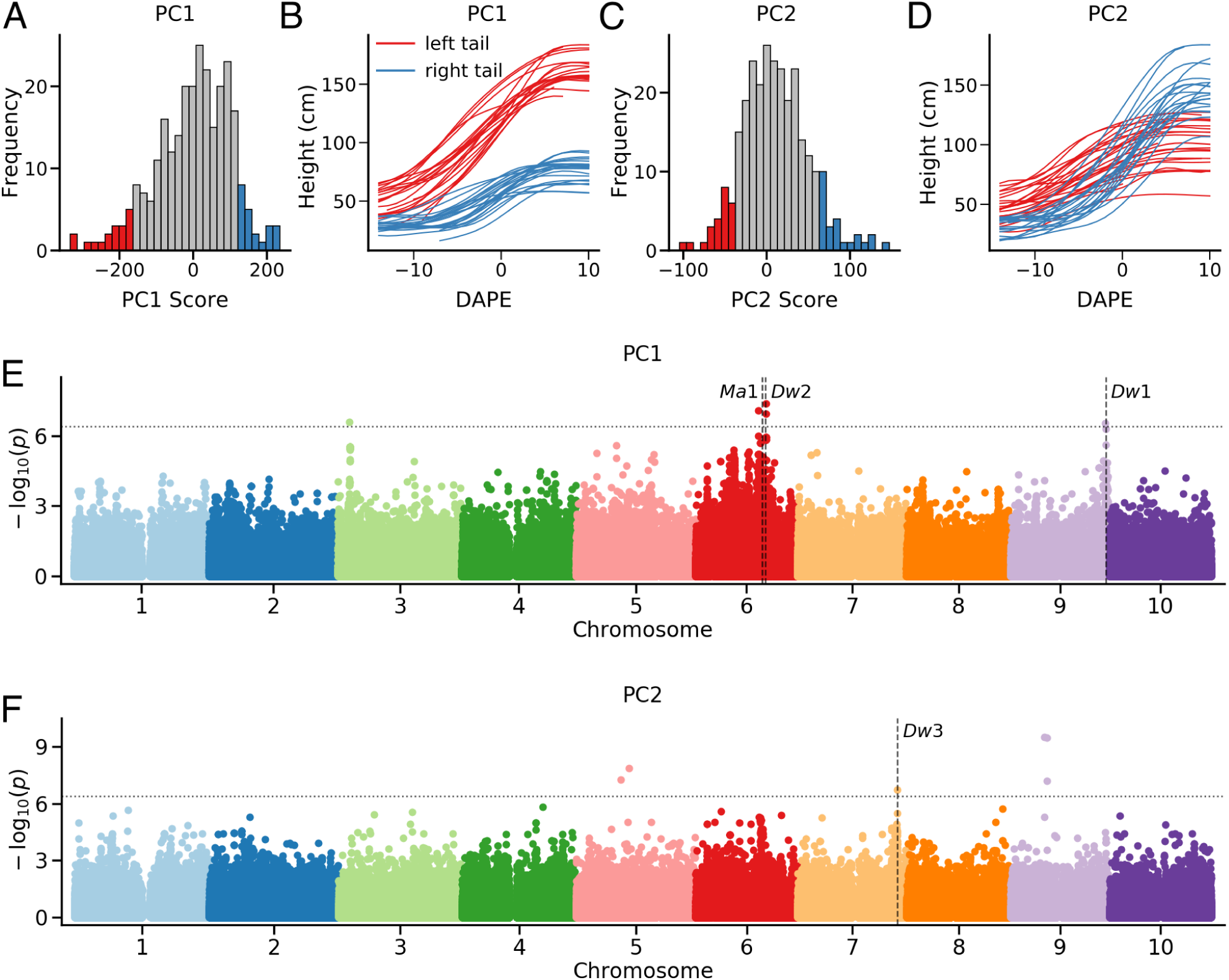
Mapping genes associated with variation in functional principal component scores among sorghum genotypes. (A) Distribution of functional principal component one scores among the 292 genotypes phenotyped as part of this study. Genotypes with the most negative values for functional principal component one are indicated in red, and genotypes with the most positive values for functional principal component one are indicated in blue. (B) Growth curves for genotypes with the most negative values for functional principal component one (red lines) and genotypes with the most positive values for functional principal component one (blue lines). (C) Distribution of functional principal component two scores among the 292 genotypes phenotyped as part of this study. Genotypes with the most negative values for functional principal component two are indicated in red, and genotypes with the most positive values for functional principal component two are indicated in blue. (D) Growth curves for genotypes with the most negative values for functional principal component two (red lines) and genotypes with the most positive values for functional principal component two (blue lines). (E) Results of conducting a genome wide association analysis for functional principal component one scores. (F) Results of conducting a genome wide association analysis for functional principal component two scores. In panels E and the positions of three cloned dwarf genes *dw1, dw2*, and *dw3* as well as the cloned maturity gene *ma1* are indicated using vertical dash lines. Horizontal dash lines indicate multiple testing corrected cutoff of a statistically significant association.

The distance between the nearest SNP significantly associated with functional principal component one and *dw1* (Sobic.009G229800) is approximately 35 kilobases. The hot region of SNPs on chromosome 6 which are significantly associated with functional principal component one spans both *dw2* (Sobic.006G067700) and *ma1* (Sobic.006G057866). The novel hit for functional principal component one on chromosome three was also identified in sequential DAPE GWAS analysis [Figure 4]. This hit is associated with a large – 17 complete genes or gene fragments – tandem array of wall-associated kinase (WAK) genes [Supplementary file]. WAKs play a role in cell elongation and expansion^43, 44^ and previous antisense experiments targeting genes in this family in arabidopsis produced dwarf phenotypes^43, 44^. The distance between the nearest SNP significantly associated with functional principal component two and *dw3* (Sobic.007G163800) is approximately 39 kilobases. The other two significant signals for functional principal component two on chromosome 5 and chromosome 9 were not associated with any immediately obvious candidate genes, which may reflect the location of these hits in low recombination/high linkage disequilibrium regions of the genome, expanding the potential distance between a trait associated genetic marker and the causal locus.

## Discussion

Engineering and computational advances are making it increasingly practical to dynamically monitor the trait values for organisms across time and development. Time-series data provides opportunities to understand the role individual genetic loci play in shaping phenotype. At the same time, extracting the greatest possible insight from time-series data requires modifications to quantitative genetic approaches traditionally employed for linking genotypic variation to phenotypic variation across an entire population at a single point in time. Above we have used time-series data from a phenotypically constrained diversity population in sorghum to evaluate a number of approaches for leveraging time-series data to identify and characterize how different loci play different roles in determining phenotype at different stages of development. A key feature which enabled our analyses was that three known large effect loci controlling height were segregating in the population of plants analyzed, yet these loci were not consistently identified in conventional single point GWAS given the phenotypically constrained set of the plants employed in the study [Figure 1]. In field studies it is often necessary to restrict the range of phenotypic diversity exhibited by an association population key traits in order to obtain meaningful and comparable trait data in a single environment^45^. Artificial selection also tends to reduce the range of phenotypic variation present within elite populations used for crop improvement^46^.

Different plants proceed through different stages of their life cycle at different rates. Comparisons across varieties must, explicitly or implicitly, employ life cycle landmarks to enable comparisons across individuals. In many field studies, traits are collected from terminal stage plants, using physiological maturity as the life cycle landmark for comparisons across individuals. In many other studies, including many greenhouse or growth chamber based ones, comparisons are performed at a fixed number of days after planting or days after germination, using planting or germination as the life cycle landmark. The sorghum association population employed in this study is samples primarily from sorghum conversion lines collected from around the world^25^. Despite the introgression of photoperiod insensitivity loci as part of the conversion process, lines vary significantly total days spent in vegetative development before transitioning to reproductive development. We experiment with using either time of planting or time of panicle emergence as the life cycle landmark for phenotypic comparisons. For these particular analyses using panicle emergence as a landmark provided significantly cleaner results with identification of known height related loci [Figure 4]. In general time-series trait data enables researchers to experiment with using different landmarks for comparisons across individuals in a population. The correct choice in any given case will depend on the specific goal of the analyses.

Simulation studies based on fitting parametric models have demonstrated that integrating longitudinal measures of phenotypes in a single population can provide increased resolution and power to identify quantitative trait loci^16^. Here we employed nonparametric models which required fewer initial assumptions about the pattern of how phenotypes change over time^47^. Like parametric models, nonparametric models can be used to accurately impute trait values at unobserved time points^24^ enabling the combined analyses of time data from larger populations given a fixed capacity in terms of number of individuals phenotyped per day. Relative to previously published functional principal component analyses, the statistical approach employed here is robust to small and uneven numbers of time point observations per sample.

Genome-wide association studies based on functional principal component weightings assigned to the time-series data from individual plants, collected on separate days, successfully identified all three known true positive genes controlling height in sorghum. Both *dw1* and *dw2* were associated with variation in the first functional principal component of sorghum height, which exhibited a consistent effect on height both before and after panicle emergence and exertion [Figure 6A, B & E]. *Dwarf3* was instead associated with the second functional principal component of sorghum height, which exhibited opposite directions of effect on height before and after panicle emergence and exertion [Figure fig:pcgwasC, D & F]. These patterns are consistent with known phenotypic consequences and molecular functions of the three known true positive genes employed in this study. *Dwarf3* and its maize ortholog *brachytic2* are involved in polar auxin transport and both reduce internode spacing^31, 48^ but showed no evidence of an influence on plant height above the flag leaf^33^. *Dwarf1* shows a strong effect on height below the flag leaf but no detectable effect on height above the flag leaf^33^. *Dwarf2* has been identified in GWAS studies using height below the flag leaf^32^ but has not been reported to influence height above the flag leaf.

In addition to successfully identifying all three known true positive height genes in sorghum, functional principal component based mapping for plant height exhibited much greater enrichment for these genes. Three statistically significant loci outside of our predefined known true positive set: a signal on the short arm of chromosome three for the first functional principal component, and two signals in the pericentromeric regions of chromosomes five and nine for the second functional principal component. Given the broader mapping intervals and slower LD decay in pericentromeric regions it was not possible to confidently conclude whether the signals on chromosomes five and nine to specific previous reports from QTL mapping or genome-wide association in sorghum. However, the chromosome three signal was validated as corresponding to a tightly mapped height QTL identified in a separate recombinant inbred line population^49^.

### Conclusions

Time-series trait data is rapidly becoming more probably available and collected in a broad range of contexts: model organisms, crops, livestock, humans, etc. Here we have demonstrated that integrating functional principal component analyses into genome-wide association studies allows the identification of known true positive genes controlling phenotypic variation in populations where these genes cannot be confidently identified from data at any single time point. In addition, we show that the association of different known true positive genes with different functional principal components describing variation in the target trait are consistent with the known biological roles and previous quantitative genetic associations of those genes. This in turn suggests that functional principal component based genome-wide association studies of time-series data can provide greater insight into the distinct roles different trait associated loci play in determining variation in a single phenotype. We employ a statistical approach to decomposing patterns of phenotypic variation over time into functional principal components which is robust to incomplete and uneven observations of different subsets of the population at different time points. This robust approach should aid the broader adoption of functional principal component based genome-wide association in a wider range of quantitative genetics contexts.

## Methods

### Plant materials and growth conditions

357 sorghum lines from SAP were planted, grown and phenotyped at (Latitude: 41.162, Longitude: -96.407) part of the the University of Nebraska-Lincoln’s Eastern Nebraska Research and Extension Center in 2016. Plant height, defined as the distance from emergence from the soil to the top of the uppermost panicle was manually scored at maturity [Supplementary file S1]. Due to the height limitation of the imaging chamber equipped in the high throughput phenotyping facility in the University of Nebraska-Lincoln’s Greenhouse Innovation Center (UNL-GIC) (Latitude: 40.83, Longitude: -96.69). Thirty-eight lines with heights in the field >2 meters were excluded from subsequent greenhouse experiments. Seeds from the remaining 319 sorghum SAP lines were sown in the greenhouse of UNL-GIC on June 15, 2016. Sorghum seeds were sown in 9.46 liter pots with Fafard germination mix supplemented with 8.8Kg 3-4 month Osmocote and 8.8Kg 5-6 month Osmocote, 1 tablespoon (15 mL) of Micromax Micronutrients, and 1800g lime per 764.5 liter (1 cubic yard) of soil. Twenty-seven sorghum lines failed to germinate or failed to grow healthily under greenhouse conditions were omitted from downstream analyses, leaving a total of 292 lines for phenotyping. Phenotyped plants were grown under a target photoperiod of 14:10 day:night with supplementary light provided by light-emitting diode (LED) growth lamps from 07:00 to 21:00 each day. The target temperature of the growth facility was between 20 − 28.3°C. After growing in the greenhouse for 40 days, all the plants were moved on to the conveyor belt which transferred each pot to the imaging chamber every two days and to the watering station each day to keep all the plants growing under a good condition. At the watering station plants were weighed once per day and watered back to a target weight, including pot, soil, carrier, and plant of 6,300 grams from July 25th to August 9th, 7,000 grams from August 9th until the termination of the experiment August 31st – 76 days after planting.

### Image data acquisition

The imaging of all phenotyped sorghum lines commenced on July 25th and continued until August 31st, 41 to 76 days after planting (DAP). Image data was collected using a high throughput phenotyping facility in UNL-GIC previously described^34^. Each plant was imaged every other day by a visible camera (piA2400-17gm, BASLER, Germany with PENTAX lens) and a hyperspectral camera (Headwall Photonics, Fitchburg, MA, USA). The visible camera was used to capture RGB images with the resolution 2,354 × 2,056 including two different zoom levels. The first zoom level was applied from 41 to 53 DAP with each pixel represented an area of approximately .45mm^2^ for objects in the range between the camera and the pot containing the plant. The second zoom level with each pixel representing an area of approximately 1.8mm^2^ was applied after 54 DAP until the termination of the experiment (76 DAP). The hyperspectral camera has a spectral range from 546 to 1,700 nm and 243 image bands. Plants were arranged so that most leaves perpendicularly face to the hyperspectral camera. A hyperspectral cube for each plant were captured at a resolution of 320 × 560 pixels. For each hyperspectral cube, a total of 243 separate intensity values were captured for each pixel spanning the range of light wavelengths between 546-1700 nm. A constant zoom level was applied for all the hyperspectral images throughout the whole experiment with each pixel represents an area of approximately 8.8 mm^2^.

### Height extraction from RGB and hyperspectral images

The estimate of plant height in RGB images uses the green index threshold method based on the equation 2 × Green /(Red + Blue)^35^. In this study, the green index threshold 1.12 was applied to separate plant pixels from other background pixels. After the whole plant segmentation was done, the number of pixels from the lowermost to the uppermost of the plant in the vertical direction were counted to represent the plant height in the pixel unit. While the stalk + panicle height from hyperspectral images were estimated using a semantic segmentation method based on the hyperspectral signatures of different sorghum organs^26^. A Linear Discriminant Analysis (LDA) model was adopted from the previous sorghum semantic segmentation project to classify each pixel to either background, leaf, stalk, and panicle^26^. After the semantic segmentation, the stalk + panicle plant height in the pixel unit was estimated by counting the number of stalk and panicle pixels on the vertical direction. Then the real plant height from RGB and hyperspectral image were obtained by using the pixel height multiply by the ratio of the real size and pixel size based on the corresponding zoom level.

### Nonparametric fitting of growth curves and missing heights imputation

The growth curve of each sorghum line was obtained by fitting the heights at different time points using the nonparametric regression. Meanwhile, the missing heights were also imputed. Let *Y*_*i j*_ be the *j*th observed phenotype of the *i*th plant, made at day *t*_*i j*_, *i* = 1, 2, …, *n, j* = 1, 2, …, *m*_*i*_, where *m*_*i*_ is the total number of days observed for the *i*th plant. To model the plant growth, we propose to use the following non-parametric model

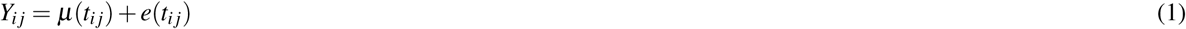

where *µ*(·) is a mean function of the phenotype development and *e*(*t*_*i j*_) is a zero-mean process associated with *i*th plant observed at *t*_*i j*_. Let *B*(*t*) = (*B*_1_, …, *B*_*K*_)^*T*^ (*t*) be a vector of B-spline basis functions, where *K* is the number of basis functions. The estimated mean function can be expressed as 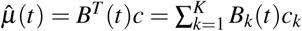, where *c* is a vector of coefficients of length *K* obtained using penalized least squares approach^24, 50^.

### Functional principal component analysis (FPCA)

The plant growth curves over time were summarized using functional principle component scores. In FPCA, the process *e*(*t*_*i j*_) in (1) is decomposed into two parts:

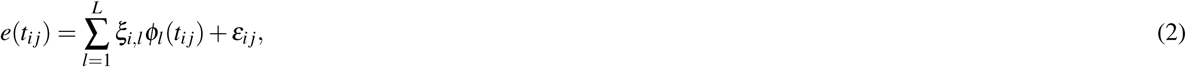

where *ξ*_*i,l*_ are zero-mean principal components scores with variance *λ*_*l*_, *ϕ*_*l*_ (*t*_*i j*_) are eigenfunctions corresponding to principal components scores, and *ε*_*i j*_ are zero-mean measurement errors with constant variance. In FPCA, eigenfunctions are orthonormal, namely 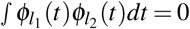, for all *l*_1_ ≠ *l*_2_ and 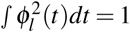, so the characteristics of phenotype development for the *i*th genotype can be represented by its principal components scores *ξ*_*i,l*_, *l* = 1, 2, …, *L*. The variance of the principal components scores, *λ*_*l*_, *l* = 1, 2, …, *L*, are sorted in decreasing order, so the first few principal components scores usually capture the majority of variation in the phenotype data. We also use B-spline bases for the approximation of eigenfunctions. The variance *λ*_*l*_ and eigenfunctions *ϕ*_*l*_ (·) are estimated by eigenvalue decomposition and the principal components scores *ξ*_*i,l*_, *l* = 1, 2, …, *L* are estimated using best linear unbiased prediction^23, 50^.

### Genotyping the SAP

DNA was extracted from seedling stage plants grown in the Beadle Center Greenhouse complex at UNL using the same seed employed for phenotyping. Sequencing libraries generated using a modified tGBS protocol^51^ were constructed and paired-end (151 bp read length) sequenced in 8 lanes of an Illumina HiSeqX instrument. This produced an average of 4 million paired end reads per sample. The quality of raw reads was controlled using the Trimmomatic/0.33 software with the parameters ‘TRAILING:20’, ‘SLIDINGWINDOW:4:20’, and ‘MINLEN:40’^52^. High quality reads were mapped to the sorghum reference genome (V4)^53^ using the BWA/0.7.17 MEM algorithm with default parameters^54^. After the alignment, the GATK/4.1 HaplotypeCaller function was used to perform SNP calling with default parameters^55^ and the raw VCF file including 358 samples was generated. Quality controls of the raw VCF including missing data rate (<0.7), minor allele frequency (MAF) (>0.01), and heterozygous rate (<0.05) were applied using customized python scripts (https://github.com/freemao/schnablelab). Missing data were then imputed using Beagle/4.1 with parameters ‘window=6600, overlap=1320’^56^. The sizes of sliding windows and the overlap windows were set to capture 10% of all SNPs on a given chromosome and 2% of all SNPs on a given chromosome, respectively. A final set of 569,305 high confidence SNPs was generated and employed in all downstream analyses.

### Genome-wide association study analyses

Three kinds of GWAS were conducted in this study: 1. Sequential GWAS of plant height based on days after planting (DAP sequential GWAS); 2. Sequential GWAS of plant height based on days after panicle emergence (DAPE sequential GWAS); 3. GWAS of the dynamic trait of the plant growth curve (FPCA GWAS).

For DAP sequential GWAS, the plant heights of 292 lines at each time point were used as the phenotypes. As different subsets of the population were phenotyped on alternating dates, height values for all plants on individual data were drawn from the values for the nonparametrically fit curves for that genotype (as described above). Genotype data of the 292 lines were drawn the original genotype data including 358 lines from SAP. After the subsetting to data from the 292 phenotyped SAP lines, any SNPs which now fell below the previous criteria used to filter SNPs for the full set of 358 lines – minor allele frequency (MAF) (>0.01) and the heterozygous rate (<0.05) – were excluded. A total of 36 GWAS were conducted from 41 DAP to 75 DAP using the same genotype data. For the DAPE sequential GWAS, X-axis (time) values for individual plant growth curves were recentered based on the date of panicle emergence. The panicle emergence was defined when the tip of the panicle is visible from the sorghum flag leaf sheath in the image. As a result, the number of lines with observed data was inconsistent across individual time points. Analyses were conducted between -14 and 10 DAPE. Within this interval, observed height values were present for at least 235 lines at each time point. For each time point, genotype information was subset and refiltered based on the set of lines with available trait data (as described above). A total of 25 GWAS were conducted from -14 to 10 DAPE using the corresponding customized marker datasets generated for phenotyped lines at each individual time point. For the FPCA GWAS, the first and second principal component (PC) scores of each sorghum line were used as trait values. The genotype data used was identical to that employed for the DAP sequential GWAS analyes.

All the GWAS were conducted using mixed linear models (MLM) implemented by GEMMA/0.95^57^. The first 5 principal components derived from the genotype data using Tassel/5.0^58^ were fitted to MLM as the fixed effect. Meanwhile, the kinship matrix calculated using “gemma -gk 1” command were fitted to MLM as the random effect. All 63 GWAS jobs were run on the Holland Computing Center’s Crane cluster at the University of Nebraska-Lincoln. The number of independent SNPs in each genotype data was determined using the GEC/0.2 software^59^. A Bonferroni corrected p-value of 0.05 calculated based on the number of independent SNPs in each specific analyses was applied as the cutoff for statistical significance in each separate GWAS analysis^60^.

## Data and code availability

Both imputed and unimputed genotype datasets used in this study have been deposited on Figshare^61^. Raw image data, including RGB, hyperspectral, fluorescence, and thermal IR images collected from each plant at each time point are being deposited and disseminated through CyVerse (**XXXXX**). The R code implementing the FPCA GWAS method used in this study has been deposited on GitHub (https://github.com/freemao/FPCA_GWAS).

## Acknowledgements

This work was supported by a University of Nebraska Agricultural Research Division seed grant to JCS and a National Science Foundation Award (OIA-1557417) to JCS. This project was completed utilizing the Holland Computing Center of the University of Nebraska, which receives support from the Nebraska Research Initiative.

## Author contributions statement

CM and JCS designed the study. CM, SL, PSS, and JCS contributed new methodology and provided datasets. CM and YX analyzed the data. CM and JCS wrote the paper. All authors reviewed and approved the manuscript.

## Supplementary materials

**Figure S1.**
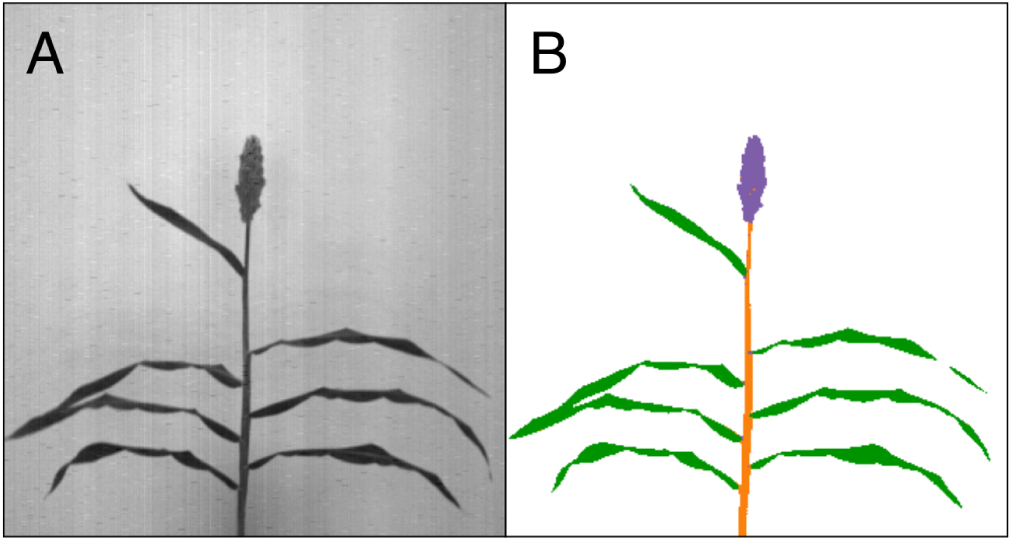
Semantic segmentation of an example sorghum plant using a hyperspectral image. (A) One of 243 distinct grayscale images necessary to summarize the information content of a hyperspectral data cube collected from imaging one sorghum plant (PI 656024) at one time point. (B) Semantically segmented image created by assigning each pixel to leaf (green), stalk (orange), panicle (purple), or background (white) classes based on intensity values for the 243 distinct wavelength intensity values recorded for that pixel within a hyperspectral data cube.

**Figure S2.**
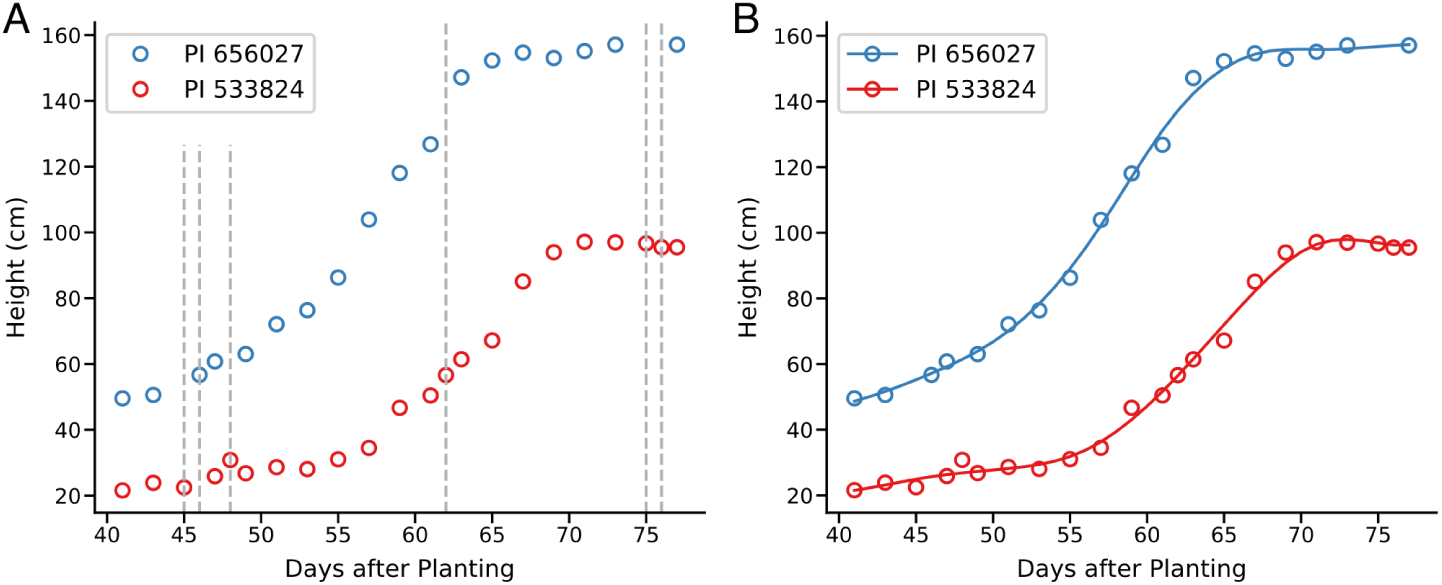
Imputing unobserved height values for sorghum genotypes through nonparametric regression. (A) Raw heights extracted from hyperspectral images for two individual sorghum genotypes (PI 656027 in blue and PI 533824 in red). Each open circle indicates a measurement extracted from an image collected on a separate day. For most time points either PI 656027 was imaged, or PI 533824 was imaged, but not both (vertical gray dashed lines) (B) Growth curves (solid lines) fit to observed values using nonparametric regression, enabling direct comparison of height values for these two genotypes at specific time points.

**Figure S3.**
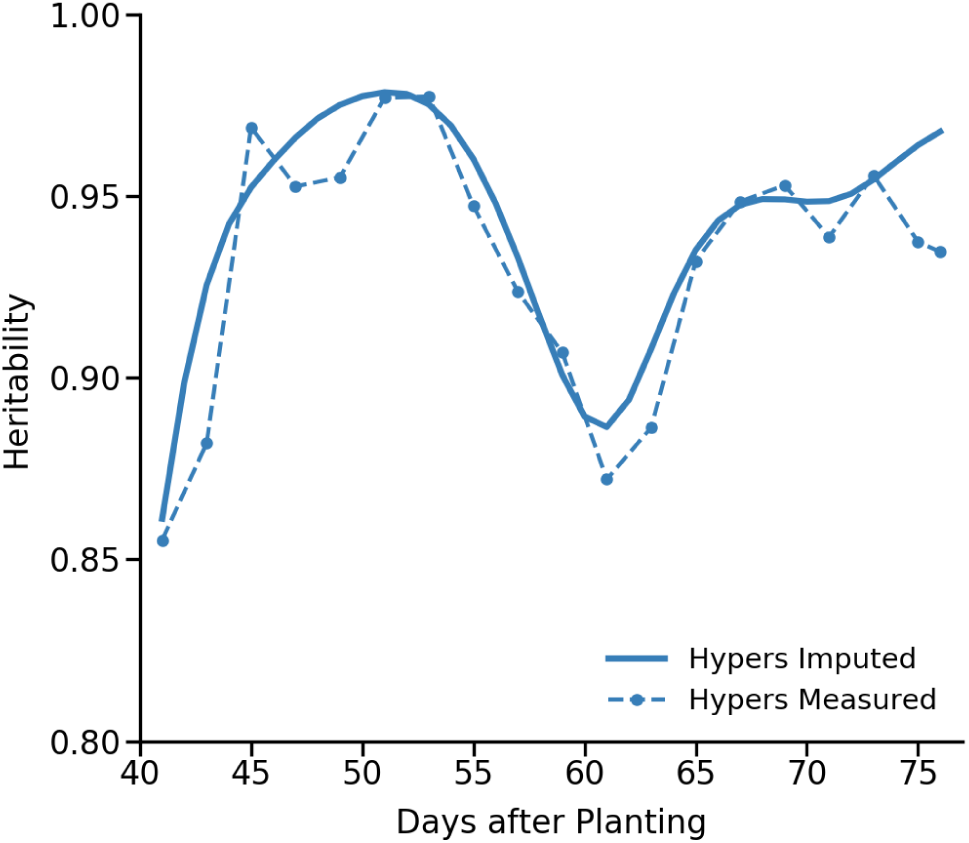
Proportion of variance explained for observed and imputed plant heights. Proportion of variance in height which could be explained by genotype for a set of twenty three sorghum lines were replicated plants were growth and imaged on the exact same set of days.

**Figure S4.**
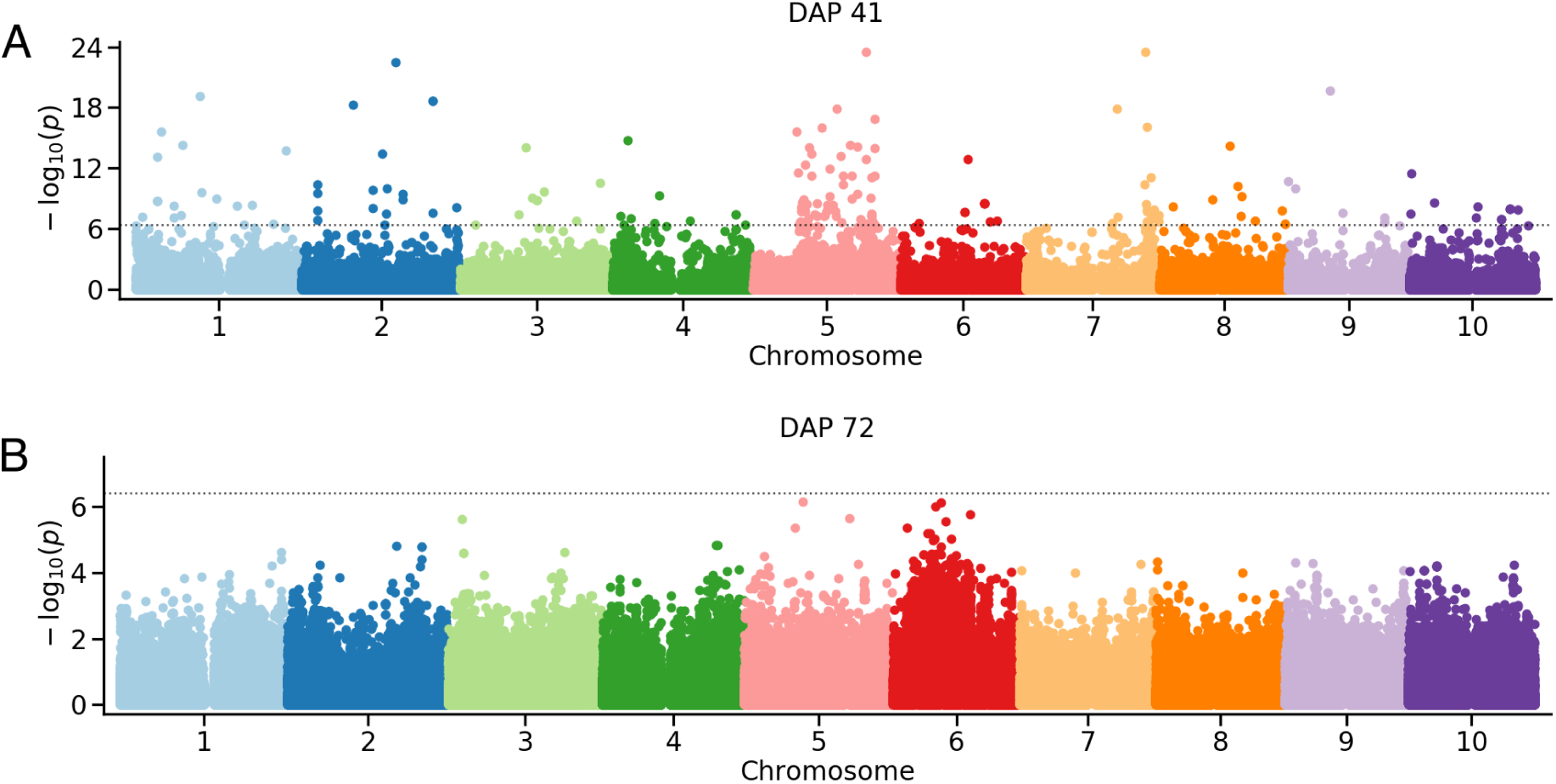
Manhattan plots for GWAS on height using height measured at 41 days after planting (Panel A) and 72 days after planting (Panel B).

**Figure S5.**
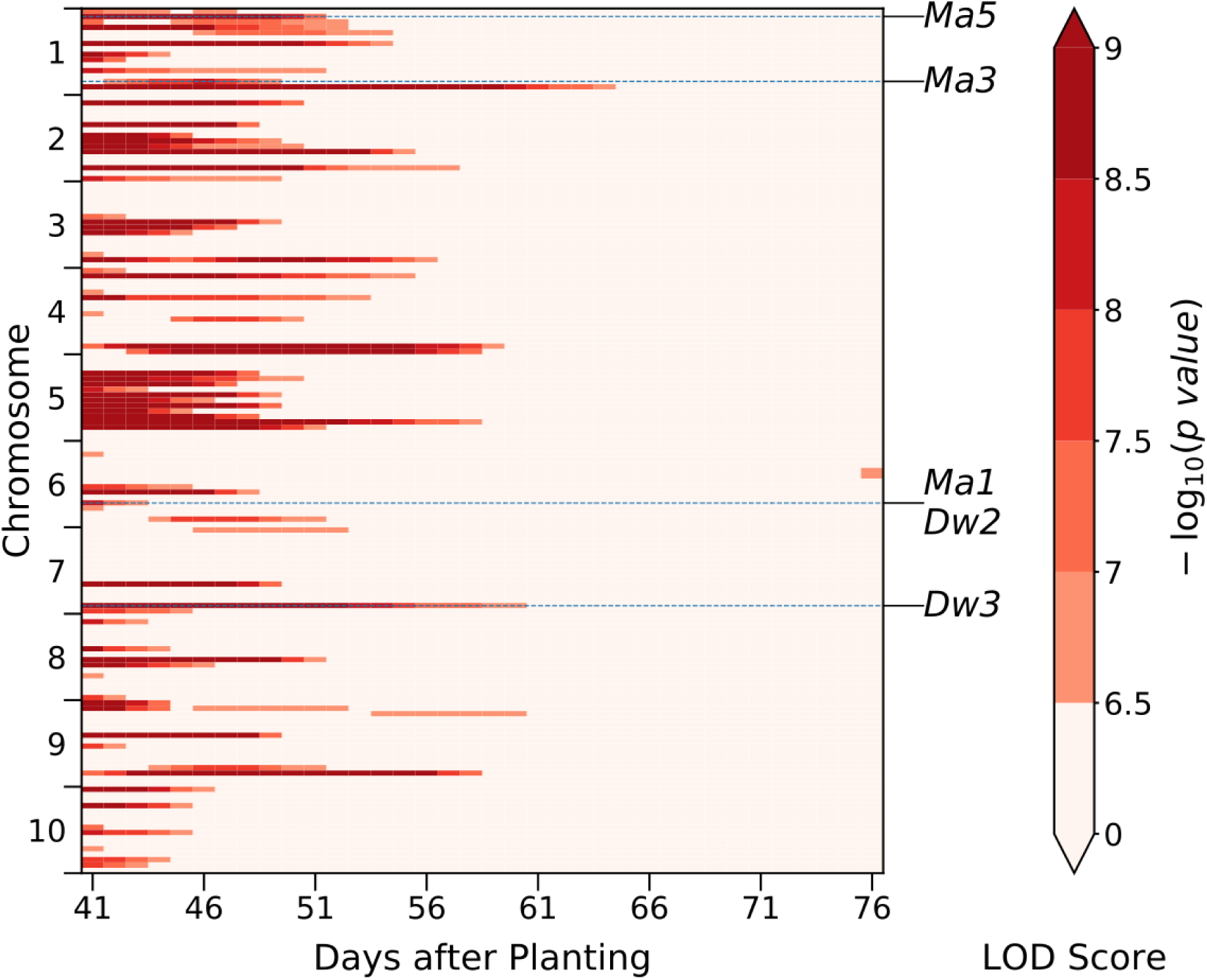
Summary of significant SNPs identified in independent GWAS for each time point anchoring on days after planting rather than days after panicle emergence. Visualization follows the detailed description provided in the legend of Figure 4.

